# Adhesion of active cytoskeletal vesicles

**DOI:** 10.1101/275115

**Authors:** R. Maan, E. Loiseau, A. R. Bausch

## Abstract

Regulation of adhesion is a ubiquitous feature of living cells, observed during processes such as motility, antigen recognition or rigidity sensing. At the molecular scale, a myriad of mechanisms are necessary to recruit and activate the essential proteins, while at the cellular scale efficient regulation of adhesion relies on the cell’s ability to adapt its global shape. To understand the role of shape remodeling during adhesion, we use a synthetic biology approach to design a minimal model, starting with a limited number of building blocks. We assemble cytoskeletal vesicles whose size, reduced volume, and cytoskeleton contractility can be independently tuned. We are able to show that these cytoskeletal vesicles can sustain strong adhesion to solid substrates only if molecular motors are able to actively remodel the actin cortex. When the cytoskeletal vesicles are deformed under hypertonic osmotic pressure, they develop a crumpled geometry with huge deformations. In the presence of molecular motors, these deformations are dynamic in nature and can compensate for an absence of excess membrane area needed for adhesion to take place. When the cytoskeletal deformations are able to compensate for lack of excess membrane area, the cytoskeletal vesicles are able to attach to the rigid glass surfaces even under strong adhesive forces. The balance of deformability and adhesion strength is identified to be key to enable cytoskeletal vesicles to adhere to solid substrates.

## Introduction

Extensive amount of work done in the past established Giant unilaminar vesicles (GUVs) as an excellent model system to study basic processes of cellular adhesion(1-5). The interactions involved in the formation of adhesion domains and the fundamental differences between the cell-cell and cell-substrate adhesion have been identified(6). Recently it has been shown that adhering vesicles act as a force generator and adhesion process itself is sufficient to induce traction forces(7). The adhesion forces can be well controlled by membrane composition of the vesicle and surface functionalization(8). During the course of strengthening of adhesion, the adhesion forces pull on the membrane, damp the membrane undulations and thus increase the membrane tension(9). Increase in adhesion strength above a critical value, causes the membrane tension to reach the critical lysis tension and consequently a bursting of vesicles is observed(10). Under conditions of specific adhesion, the lateral forces from the surface come from the attraction between the receptors in membrane and the ligands on the surface. In going from an unbound state to a bound state, the vesicles make significant shape transformation(9-12). This adhesion induced shape transformation have been successfully explained by minimizing the free energy in the framework of the Helfrich theory of elastic cells(13, 14). So far, the insight into adhesion process through the model system lacks involvement of the cytoskeletal coupling to the membrane.

Here we elucidate the role of the presence of a cytoskeletal cortex on the adhesion process. We show that a strong coupling of the actin network to the membrane causes dampening of the membrane fluctuations and an increase in membrane tension(15, 16). When a tensed cytoskeletal vesicle binds specifically to a glass surface using biotin streptavidin as ligand-receptor pair the membrane tension increases beyond the lysis tension. This increase in membrane tension is due to the need of the vesicle to deform to gain adhesion area, which in turn requires excess membrane area. However, the excess membrane area is already used up by the cortex binding and hence is not available for the vesicle to gain adhesion energy. In the absence of free excess membrane area, the receptor binding is sufficiently strong to tense the membrane beyond the critical lysis tension. We observe that for a given ligand-receptor density for which the cortex free vesicles show strong adhesion, the cytoskeletal vesicles open pores and burst.

Our work pin points the crucial dependency of adhesion on the availability of excess membrane area. Since cytoskeletal to membrane coupling is opposing the vesicle deformation to gain adhesion, we provide the cytoskeletal vesicles some excess membrane area by applying additional hypertonic stress. Under the reduced volume condition, the vesicles can then develop large deformations. Our experiments show that only active deformations of the cytoskeleton can provide the excess membrane area required to gain adhesion. Therefore, adhesion of a cytoskeletal vesicles relies on the ability of the vesicle to actively remodel its cytoskeleton. The weakly adhered active cytoskeletal vesicles make a transition from weak adhesion regime to strong adhesion regime under hypertonic osmotic stress, without rupturing their membrane.

## Materials & Methods

### Reagents

Egg L-α-phosphatidylcholine (Egg PC) lipids were ordered from Sigma (P3556) in powder form and dissolved at 50 mg/ml in a chloroform/methanol mixture (9:1, v/v). 1,2-dioleoyl-*sn*- glycero-3-[(*N*-(5-amino-1-carboxypentyl)iminodiacetic acid)succinyl] (nickel salt) lipids (Ni-NTA); 1,2-distearoyl-*sn*-glycero-3-phosphoethanolamine-N-biotinyl(polyethylene glycol)- 2000] (ammonium salt) were ordered from Avanti Polar Lipids. The mineral oil was from Sigma-Aldrich (M3516) and the silicone oil (viscosity 50 cSt) was from Roth (4020.1). Decane was from Sigma-Aldrich (D901). Biotin BSA (A8549) and streptavidin were also purchased from Sigma Aldrich.

### Proteins

Proteins were purified according to previously published protocols(17). G-actin and muscle myosin II were from rabbit skeletal muscle. The fragment of *Xenopus laevis* anillin spanning amino acids 1 to 428, excluding the myosin binding site, was cloned into a pET-28a vector and purified from *Escherichia coli*, with His tags on both termini.

### Buffer solutions

We mixed the solution to be encapsulated on ice immediately before vesicle production. Anillin, myosin II, and G-actin were added to a polymerization buffer (pH 7.2). The chemical composition of the solution (including salts from protein buffers) consisted of 10 mM imidazole, 1mM MgCl_2_, 1mM adenosine triphosphate (ATP), 1mM EGTA, 30mM KCl, 2mM dithiothreitol, 300mM sucrose, 0.5 μM Alexa Fluor 488 phalloidin. The outside solution for production of vesicles was made of glucose, whose osmotic pressure was adjusted 10 to 15mosmol higher than the inside solution.

### Vesicle production

Vesicles were produced using the cDICE method(16, 18). Briefly, this method consists of a cylindrical rotating chamber successively filled with a glucose solution to collect the vesicles, a lipid-in-oil solution to saturate the oil/water (O/W) interfaces, and decane as the continuous phase in which droplets were produced. The protocol to disperse the lipids in the oil solution has been published elsewhere(16). The solution containing the cytoskeletal elements was injected from a glass capillary tube by inserting the capillary’s tip in the decane. Because of centrifugal force, droplets detached from the tip. The droplets then moved through the lipid-in-oil solution where they were coated by first one lipid monolayer and then by a second lipid monolayer while crossing the O/W interface. The two monolayers zipped together to form a bilayer. Vesicles were collected in the glucose solution, which was sucked with a micropipette once the chamber was stopped. For the process to succeed, the osmolarity of the encapsulated solution has to match that of the glucose solution. The whole process was completed in a cold room maintained at 5°C to prevent fast polymerization of the cytoskeleton. We produced vesicles in a span of 2 min. Although cDICE is a high-yield method, resulting in hundreds of vesicles under most conditions, encapsulating proteins at high concentrations (10μM of actin and up to 1.5μM of anillin) resulted in a decrease of the yield. At the highest protein concentrations, a 100μl sample contained about 50 vesicles.

### Adhesion protocol

BSA-Biotin and streptavidin were used to coat the coverslips to make the vesicle adhere. The stocks and working solutions of BSA, BSA-Biotin and Streptavidin were all prepared in 1x PBS. To functionalize the coverslips, they were first incubated for 20 minutes at room temperature with mix of 1mg/ml BSA-Biotin and 1mg/ml BSA followed by a couple of washing with 1x PBS and then further incubation with 0.5mg/ml streptavidin. PEG-Biotin lipids in the membrane were kept 1% for the strong adhesion. 5μM KCL was added in the vesicle suspension to make the membrane-embedded biotin, bind to the functionalized glass surface.

In addition to lowering the percentage ratio of BSA-Biotin and BSA, we also added 0.5% PEG2000 into the vesicle membrane to lower the adhesion strength between the coverslip and the membrane.

### Deswelling protocol

Vesicles were deswelled by adjusting the surrounding osmotic pressure in a diffusion chamber. The chamber consists of two compartments made of flat o-rings (20 mm in diameter and 2 mm thick) and separated by a membrane (Merck Millipore) with a pore size of 0.22μm. First, at t=0, a 600mOsm solution was added on top of a permeable membrane, and after about one hour a 833mOsmo solution was added.

The vesicles are confined in the bottom compartment and their surrounding osmotic pressure is changed by adding a glucose buffer on the top compartment. The osmotic pressure equilibrates in the chamber via the diffusion of the glucose through the polycarbonate membrane (pore size: 0.22μm) which separates the two compartments. The kinetics of the increase of the osmotic pressure in the bottom compartment is computed from the measurement of the osmotic pressure on the top compartment (See Fig. S1).

### Imaging and analysis

Vesicles were imaged with a Leica Microscope DMI3000 B and a 63× numerical aperture (N.A.) 1.3 oil immersion objective for bright-field microscopy and epifluorescence, in combination with a Hamamatsu ORCA-ER camera.

Confocal imaging was acquired using Leica TSC SP5 and a 63x N.A. 1.4 oil immersion objective. The 3D reconstruction using the confocal stack was done using software Imaris.

## Results and Discussion

### Cytoskeletal vesicles

Our model system is a giant unilamillar vesicle (GUV) that has a crosslinked actin cortex anchored to its inner leaflet. The His-tagged anilin is responsible for both crosslinking the actin and coupling the actin network to the Ni-NTA (nitrilotriacetic acid) lipids that are incorporated into the membrane. We call the vesicle that has an actin cortex a cytoskeletal vesicle (Fig. 1(a)). Protein encapsulation occurs during vesicle preparation using the cDICE method adapted for this system(16). By adding myosin motors to the network, we add contractility and hence activity. Depending on presence or absence of motor proteins, we call cytoskeletal vesicles active vesicles or passive vesicles, respectively. The actin cortex is formed by encapsulating 10μM of G-actin and 1.5μM of anilin at 4°C. We induced contractility into the cortex by adding an additional 0.1μM of myosin motors into the protein mix.

**Fig. 1:**
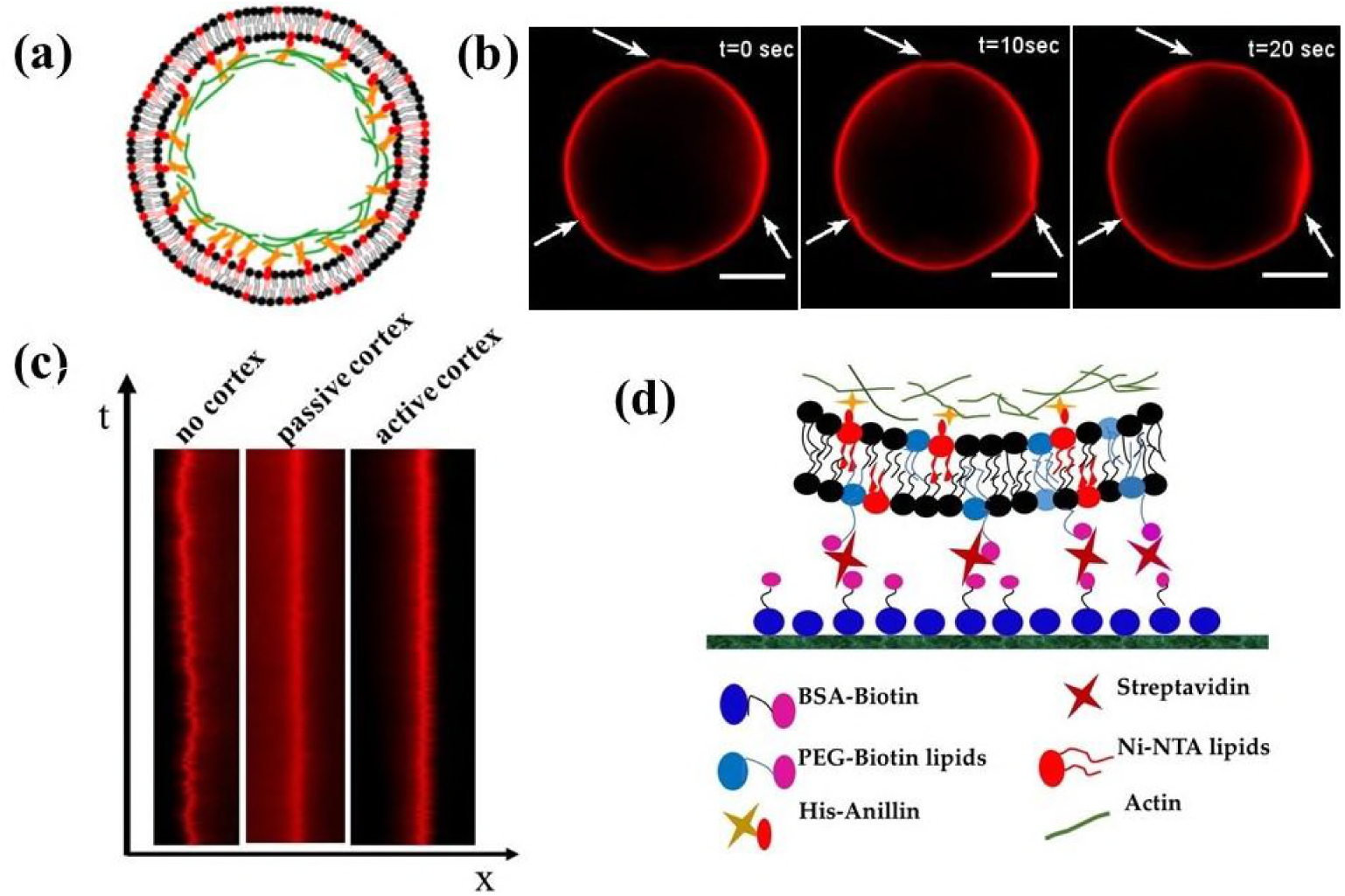
(a) The model system consists of GUVs with 10% Ni-NTA, PEG and PEG-Biotin lipids incorporated in it. Actin bundles are bound to membrane through the His-tagged Anillin, the actin crosslinker. (b) The vesicles show random changes in shape caused by the dynamic nature of the actin network. Here, the membrane is labeled with Texas Red (c) Kymographs showing loss of membrane undulation in the bilayer due to membrane-cortex adhesion. (d) The vesicles are bound to

The passive vesicles maintain their spherical shape in suspension, while the active ones change their shape with time (Suppl. movie S1) due to the active stress generated in the cortex by the myosin motors. The characteristic time scale over which we observed the random shape changes is much larger than the time scale of membrane undulations. The active cortex pushes and pulls on the lipid bilayer (suppl. movie, S1) causing tiny vertices to appear on the vesicle surface (Fig. 1(b)). We did not observe any membrane undulations such as we saw in the cortex free vesicles in the cytoskeletal vesicles (fig. 1(c)). This is due to the anchoring of the shear elastic actin cortex to the membrane, which kills the fluctuation modes. These observations already indicate that cytoskeletal vesicles have higher membrane tension than the cortex free vesicles.

### Adhesion of the cytoskeletal vesicles

The specific adhesion strength between the vesicle and the attractive glass surface can be controlled by tuning the Ligand-receptor density between the two. We used Biotin-streptavidin as ligand-receptor pair to make vesicles adhere to the glass (Fig. 1d). We used two different ligand densities at the glass surface by coating the glass with BSA-Biotin and BSA mixed at two different volume ratios, 70:30 and 50:50. The membrane was doped with 1% PEG-Biotin lipid to make the vesicles bind to the glass specifically. We begin with the adhesion of cortex-free vesicles. We observed that for both ligand densities the cortex-free vesicles bind to the rigid glass surface and adopt spherical cap shapes (Fig. 3(a)). Some of the vesicles gets leaky but maintain their shape. These observations are consistent with previously published work which showed that a vesicle with constant volume adopts a spherical cap shape under strong adhesion conditions(8). By contrast, the cytoskeletal vesicles were observed to burst in the strong adhesion regime. On a glass surface coated with a mix of 70% BSA-Biotin and 30% BSA, all the cytoskeletal vesicles burst within a minute of touching the glass surface. Lowering the adhesion strength (50% and 50%), we observed that some size selection occurs and cytoskeletal vesicles with diameter smaller than approximately 20μm are stable for more than 30 min. Our observations show that the surviving vesicles forms contact area with diameter ≤ 14.4 ±1.9 μm. Bigger vesicles form larger contact area before they burst. A collapsed actin network and supported bilayer membrane is observed at the site where a large vesicle is bursted by the strong adhesive forces. One possible reason for the observed size dependency is the local curvature of the vesicle at the contact area, which defines accessibility of the binding partners. Therefore, the elasticity of the cortex limits the spreading dynamics(19). The large contact area of the large vesicles (diameters >20μm) cause large lateral forces which increase the membrane tension beyond the lysis limit, causing the cytoskeletal vesicles to burst.

**Fig. 2:**
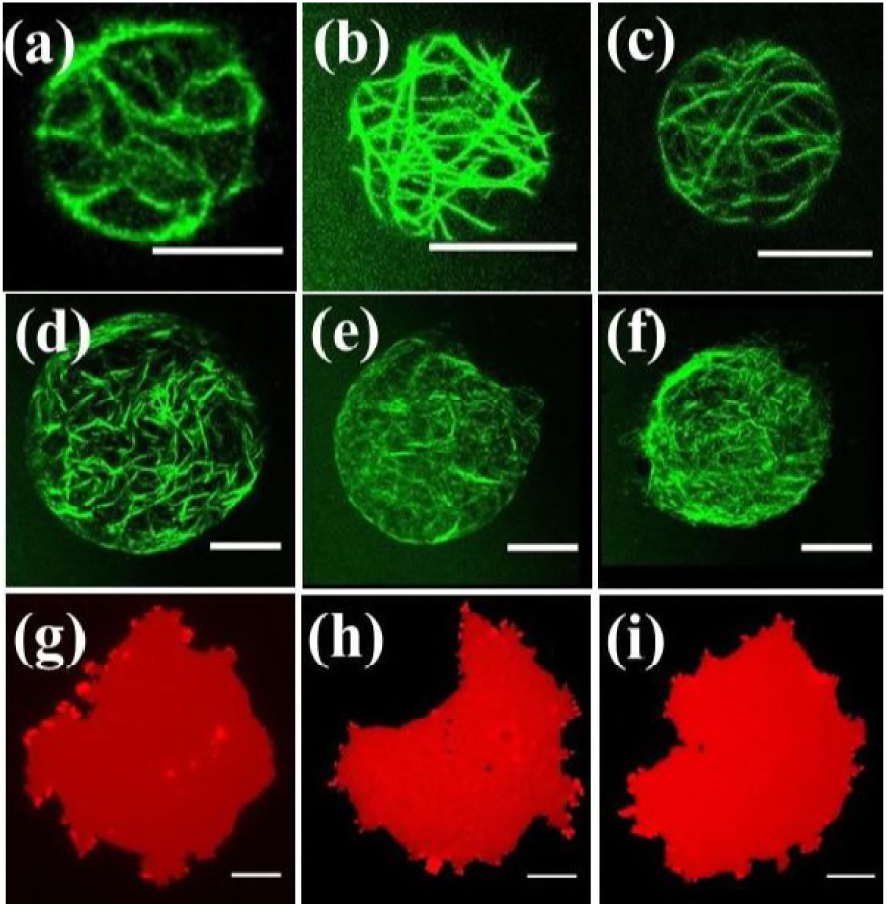
Cytoskeletal vesicles with diameters less than 20μm survive strong adhesion. (a—c) Adhesion patches corresponding to three individual cytoskeletal vesicles that survived strong adhesion. No larger than 15μM contact diameter was observed for strong adhesion. Contact diameter more than 15μm results in bursting of vesicles. (d—f) The vesicles that burst show actin fibers attached to the surface in the contact area, giving an estimate of the contact area achieved before bursting. (g—i) The membrane of the bursting cytoskeletal vesicles form a supported bilayer on the functionalized glass surface. The area of this supported bilayer was consistently 2—3 fold bigger than the actin patch in the contact area of the burst vesicle. The scale bar here is 10μm.

**Fig. 3:**
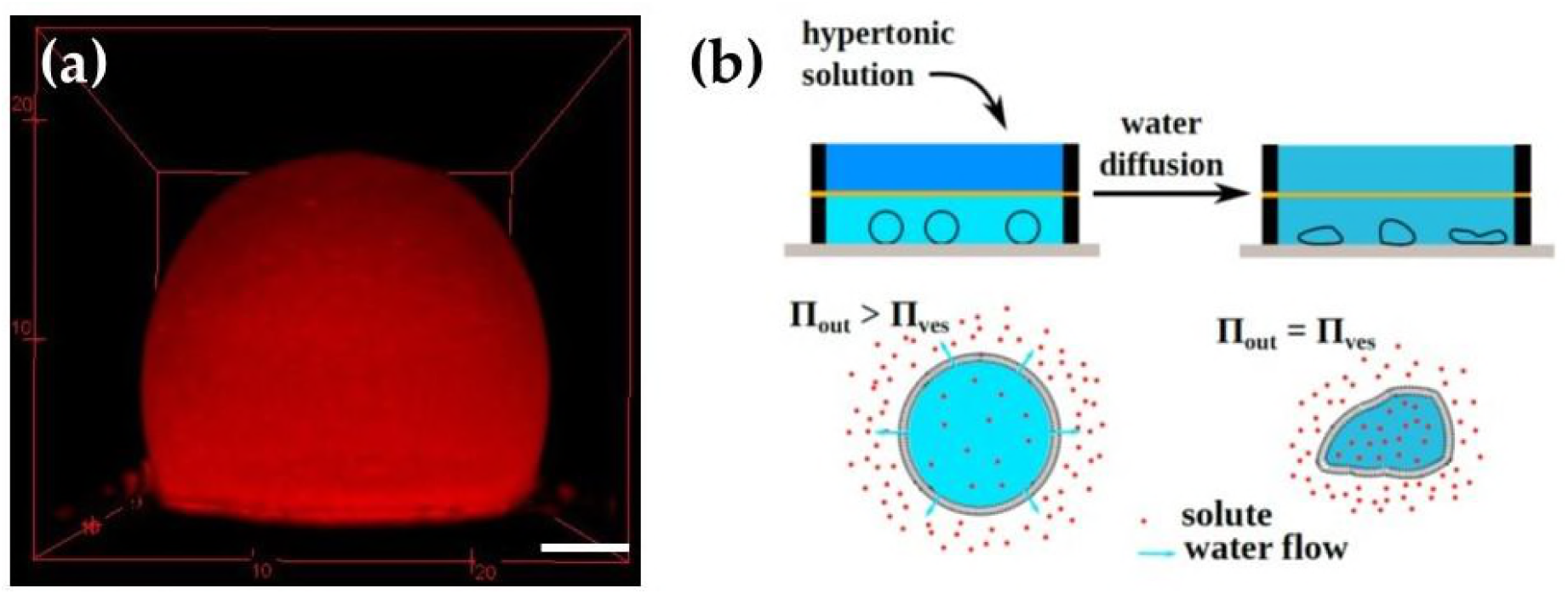
(a) A strongly adhered cortex-free vesicle adopts a spherical cap shape in its equilibrium bound state. (b) The two-level chamber applies hyperosmotic stress to the vesicle. The top chamber is filled with a glucose solution that has higher osmotic pressure than the glucose solution in the bottom chamber. We selected a difference in the osmotic chamber that would produce a reduction in vesicle volume of 40%. The two chambers were separated by a membrane that had

The finding that the cytoskeletal vesicles burst, while cortex-free vesicles are able to adhere at a similar ligand-receptor density can be attributed to the difference in availability of excess membrane. Cortex coupling has already been reported to limit available excess area and thus to limit the vesicle’s ability to form membrane tubes under hydrodynamic flow(20). To gain in adhesion a vesicle needs to deform and deformations are possible only at the cost of membrane undulations and thus by increasing the membrane tension. Membrane tension limits the gain in adhesion once the excess membrane area has been consumed. Since the membrane undulations are missing, owing to the coupling of membrane to the cortex, the membrane is in tensed state in case of the cytoskeletal vesicles. Therefore, for the cytoskeletal vesicle the increase in contact area beyond the cut off causes lysis of the membrane. In a next series of experiments, we aimed to increase the available excess area by hypertonic osmotic stress in order to enable the strong adhesion of cytoskeletal vesicles.

### 2.3 Deformations in cytoskeletal vesicles under hypertonic osmotic stress

Since the limiting parameter for creating adhesion is a lack of excess membrane area, we applied a hypertonic osmotic stress using a two-level diffusion chamber (Fig. 3b) to deflate the vesicles to a reduced volume of ν=0.6 (40% volume loss). The reduced volume is defined by the ratio between the volume of liquid present in the deformed vesicle and the volume enclosed by a sphere with the same surface area. We compare the deformations of passive and active vesicles in the non-adhering state under hyper osmotic pressure to pinpoint the effect of myosin motors on shape adaptation.

Cortex free vesicles show the well-described morphological deformations predicted by the minimization of curvature energy of the lipid membrane(21-23) (fig. 4a, suppl. Movie S2). In contrast, the passive cytoskeletal vesicles remain mostly spherical for up to a 10% increase in the osmotic pressure without changing their volume (Fig. 4b,c). Since the volume of the passive vesicle does not change significantly, we estimate that the resulting applied pressure reaches 0.11 atm. The reduced volume and the vesicle shape remain almost the same as the surrounding osmotic pressure increases from 450 to 500mOsm (Fig. 4b,c). Increasing the osmotic pressure further leads to a further compressive stress build up and, finally, to an abrupt deformed shape change (∼6% radius decrease). After this abrupt deformation, when the pressure reaches 520mOsm, the cortex stability again resists further deformations, until the pressure exceeds 540mOsm (Fig. 4c) and a second sudden shape change occurs. After the second sudden event of cortex shape change, the radius of the vesicle starts decreasing monotonically as external pressure increases.

**Fig. 4:**
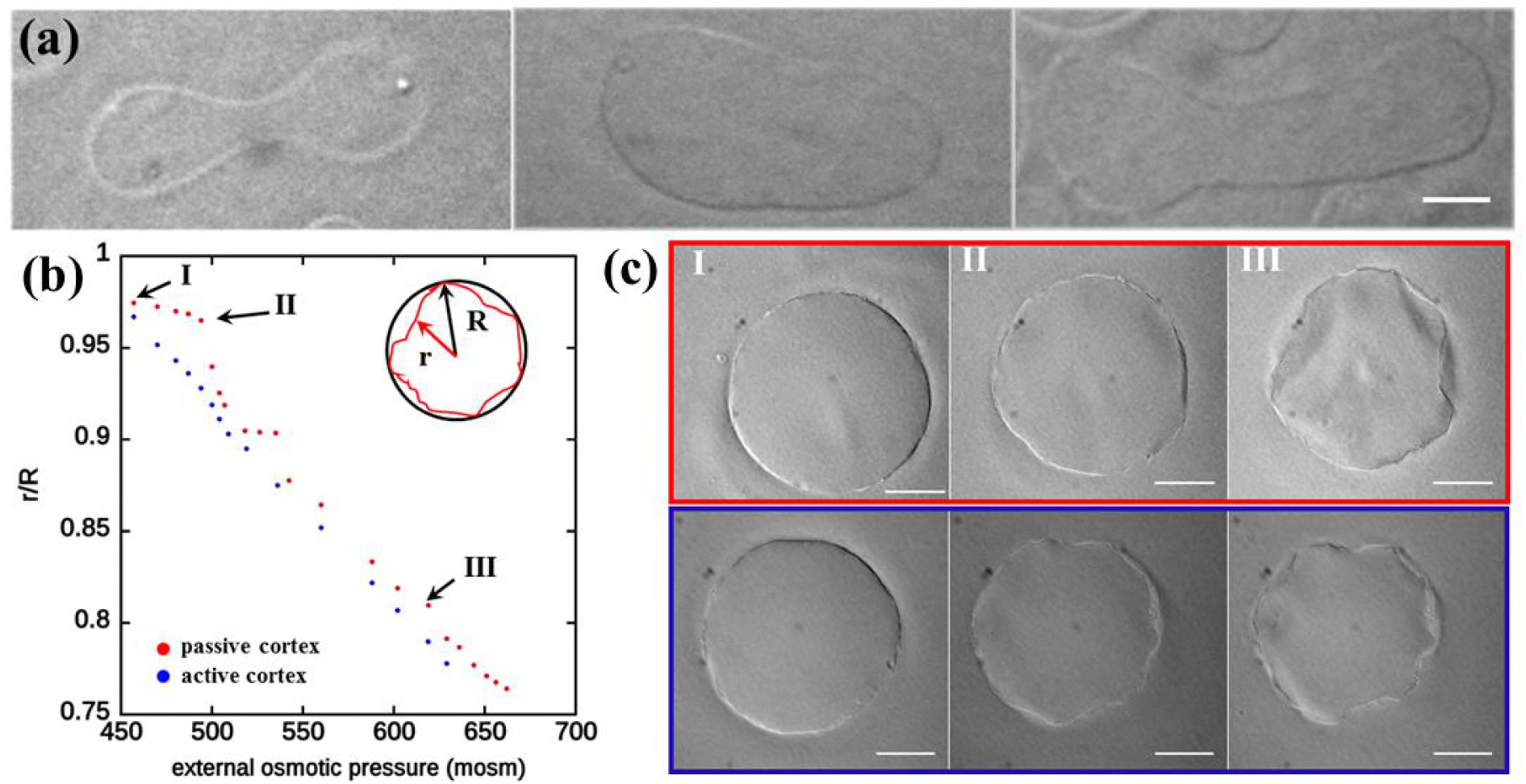
(a) Deswelling of the cortex free vesicles shows well-documented shape transformations. (b) From point I to II, the change in external osmotic pressure does not cause any shape remodeling for the passive vesicles but causes continuous shape remodeling for active vesicles. (c)All three images taken from the plot in (b) show similar shapes after remodeling, the only difference being the mode of reaching that shape. For passive vesicles the mode is abrupt and for active it is continuous.

In contrast to these discontinuous deformations of the passive cytoskeletal vesicles, the presence of 0.1μM of myosin motors enable vesicles to adapt continuously to the osmotic pressure change. Indeed, the myosin contractile activity pulls at the membrane and the limiting parameter to deformation is now the membrane tension of the vesicle. We observe a continuous remodeling of vesicle shape, without any sudden instabilities (Fig. 4b,c). The final equilibrium shape of both passive and active vesicles is comparable (Suppl. Movie S3). Both types show a complex morphology with many edges, resembling a crumbling transition of an elastic shell (Fig. 4c at point III (Suppl. fig. S2)), as predicted for spherical elastic shells submitted to a constant compressive rate(24).

### Adhesion of cytoskeletal vesicles under hypertonic osmotic stress

Since adhesion relies on the availability of excess area, we tested the adhesion process of active and passive cytoskeletal vesicles having diameters larger than 20μm under hypertonic stress. To avoid immediate bursting of vesicles having diameters larger than 20μm, we reduced the adhesion strength between the membrane and the glass by adding 0.5% PEG- 2000 lipids to the membrane and by coating the glass with a mix of 35% Biotin-BSA and 65% BSA instead of a 50—50 mix. Cytoskeletal vesicles are still unstable and we observed the bursting of the vesicles even under the lower adhesion strength. Due to a lower density of adhesion molecules and the presence of PEG lipids in the membrane, the formation of contact area slows down and rupture is delayed by around 10-15min. Since the assembly of the diffusion chamber takes just about 1—2 minutes, the cytoskeletal vesicles are under hypertonic stress long before the lysis point. To estimate the contact angle at the initial stage, when the osmotic pressure is beginning to build, we performed control experiments. In the control experiments we acquired the z-stack of the vesicles in the first 10 min of the adhesion process under no osmotic stress. We imaged 20 such vesicles and Fig. 5(a) shows an example of the shape of a cytoskeletal vesicle in the initial bound state. We found no difference in the geometry of the initial bound state between the active and passive vesicles and the average contact area was around 122 °. We chose the glucose concentration in the top chamber to give an equilibrium osmotic pressure that would reduce the vesicle volume by around 48%. In 70 minutes the solutions in the two compartments reached equilibrium (calibration plot in supplementary Fig. S1) and we began imaging.

**Fig. 5:**
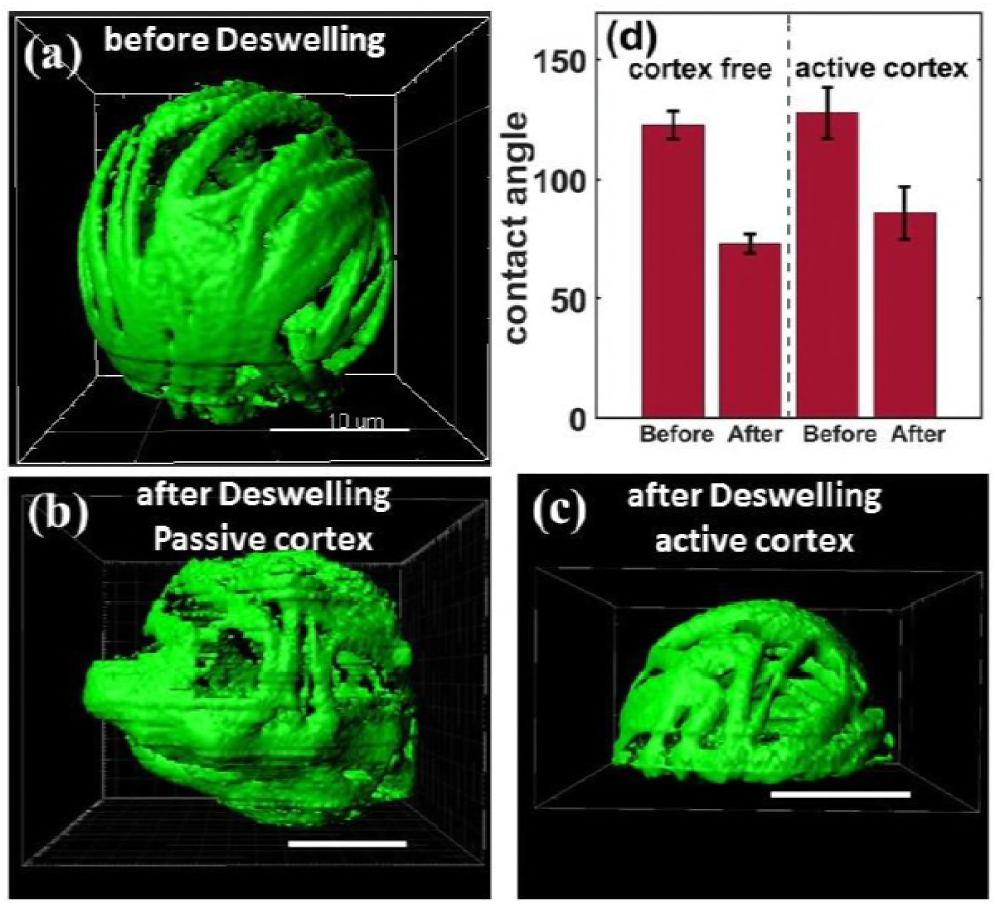
(a) A cytoskeletal vesicle (active/passive) shows weak adhesion before deswelling with an average contact area of around 122 °. (b) The passive vesicles show random shape remodeling and no increase in adhesion strength for most of the vesicles. (c) The active vesicles after deswelling gain in contact area and show a decrease in the contact angle. (d) The almost equal change in contact angle from the non deswelled to the deswelled state of the weakly adhered cortex in the free and the active vesicles shows the availability of the excess area in deformations for gain in adhesion strength. 20 vesicles were imaged to estimate the contact angle and the error bars show ±SD.

We observed that the deflated cytoskeletal vesicles remain stably adhered to glass even after 2hrs of contact with the glass without any observable bursting event. We observed a striking cortex-activity-dependent shape transformations in cytoskeletal vesicles under hypertonic stress. After a volume reduction of around 48%, passive vesicles became strongly and irregularly deformed (fig. 5(b)) while all active vesicles adopted the shape of a smooth spherical cap (fig. 5(c)).

An increase of adhesion area of the active cytoskeletal vesicles suggests that the excess membrane area created by developing deformations in the vesicles is effectively used to gain adhesion strength. Thus myosin activity is essential for the cortex remodelling in order to gain adhesion area. Passive ones lack the ability to increase adhesion area and just crumple under the osmotic pressure change. Resulting forms of the passive vesicles are indistinguishable in non-adhered and adhered conditions, which demonstrates that adhesive forces alone are not sufficient to induce a shape change of the elastic shell. For both cortex free vesicles and active cytoskeletal vesicles, the contact angle changes from about 122o in weak adhesion conditions to about 70 ° in strong adhesion conditions, as shown in Fig. 5(d). The contact angle for passive vesicles could not be determined after deswelling due to its highly deformed random shape near the surface. A comparison between the final shape gained by the passive and active vesicle can be seen in supplementary movie S4.

The passive vesicles show abrupt changes in shape caused by a sudden buckling of the actin cortex. For non-adhering cytoskeletal vesicles, we observed that the cortex crumples only when the osmotically induced deformation forces are sufficiently high (Fig. 4(b)). In comparison, the attractive forces of the small adhesion zone are too small to induce any crumpling events. Consequently, the excess area created remains within random locations of the deformations and is not available to change the adhesion area.

On the other hand, the presence of myosin motors induces activity in the cortex and makes it easy for the cortex to adapt. It is now an actively remodeling cortex and can adapt easily to external forces. The osmotic pressure and the adhesion forces both are able to induce deformations. During adhesion- area formation, the osmotic pressure continuously yields sufficient excess area which is then continuously pulled laterally by the adhesive molecules. Excess area of the osmotically induced deformations is thus made available for adhesion by the motor activity. Our experiments show that not the presence of excess membrane area but its availability is the factor that helps the vesicle gain in adhesion. The availability of excess membrane area depends on the ability of the actin cortex to remodel actively.

## Conclusion

Our results describe the very basic and essential process of balancing cortex attachments and adhesion induced contact area formation. By altering the degree to which the actomyosin cortex is anchored to the membrane and by reducing the volume of the vesicle, we can explore the complex interplay between membrane tension, cytoskeleton elasticity, and active forces in the context of cell adhesion. Upon adhesion to a rigid substrate, cytoskeletal vesicles need to accommodate the shape of the coupled cytoskeleton/membrane shear elastic material. While the lipid membrane is fluid and non-stretchable, the elastic cytoskeleton can be sheared and stretched. These two mechanical properties result in a strong constraint for the system. Cystoskeletal vesicles can withstand strong adhesion provided that two conditions are fulfilled. First, some excess membrane area must be available to allow shape remodeling without overcoming the critical lysis membrane tension; this is evidenced by the fact that large passive vesicles with the surface to volume ratio of a sphere burst when starting to adhere. Second, the actin cortex should be able to undergo remodeling to accommodate the substrate configuration. Under the experimental conditions of our study, molecular motors are required to generate active forces to dynamically remodel the cytoskeleton. In the case of a passive cortex, its stiffness prevents the vesicle from spreading and adhering on a substrate. This study provides a conceptual framework for other research addressing complex questions about cell adhesion. For example, it would be interesting to investigate proteins that exhibit adhesion that has a finite lifetime in order to better understand recruitment and formation of adhesion patches that drive the dynamics of adhesion.

## Acknowledgments

Research was supported by ERC-SelfOrg and partly by the SFB863 and the Nanosystems Initiative Munich.

